# Singing-related reorganization of the song motor circuit is accompanied by a shift in hormonal modulation of song

**DOI:** 10.1101/2020.08.31.274100

**Authors:** Mariana D. Rocha, Jes Dreier, Jonathan Brewer, Manfred Gahr, Michiel Vellema

## Abstract

Sex hormones are essential modulators of birdsong. Testosterone, and its active androgenic and estrogenic metabolites, 5α-dihydrotestosterone (DHT) and estradiol, can re-shape the brain circuits responsible for song learning and production. The differential mechanisms of action of these different hormones during song development and song maintenance are, nonetheless, not fully understood. Here we demonstrate that unlike testosterone, DHT treatment does not induce singing behavior in naïve adult female canaries that have never previously produced song. However, in birds with previous testosterone-induced singing experience, DHT alone is enough to promote the re-acquisition of high quality songs, even after months of silence. In addition, we show that the synaptic reorganization that accompanies vocal motor skill development requires more than DHT-induced androgen receptor activation. These results indicate that vocal motor practice will persistently modify the hormone-sensitive brain circuit responsible for song production, suggesting a mechanistic differentiation in the hormone-dependent regulation of the initial vocal motor skill acquisition and later re-acquisition.

## Introduction

Sex hormones are critical modulators of adult neuroplasticity and behavior. Sex hormones can either be produced locally in the brain or be synthesized in the gonads and reach the brain through blood circulation (Diotel et al., 2018). Additionally, circulating testosterone can be converted in the brain into its active metabolites through enzymatic action: into DHT (5α-dihydrotestosterone) by 5α-reductase, or into estradiol (17β-estradiol) via the enzyme aromatase. Through their interactions with androgen and estrogen receptors in the brain, sex hormones can mold brain circuits, and thus shape animal behavior in complex ways (Chen et al., 2013; Diotel et al., 2018; Gahr, 2004; Galea, 2008).

Birdsong is a complex motor behavior produced as a result of self-reinforced vocal imitation and exhibits many parallels to human speech (Bolhuis et al., 2010). Birdsong is known to be modulated by sex hormones in the song system, the network of interconnected brain nuclei responsible for controlling song learning and production. Seasonal changes in the vocal performance of songbirds are highly correlated with changes in testosterone levels (Nottebohm et al., 1987; Smith et al., 1997; Tramontin and Brenowitz, 2000; Voigt and Leitner, 2008). These changes in testosterone levels are accompanied by gross anatomical and cytoarchitectural restructuring of the song system (Balthazart et al., 2008; Gahr, 1990b; Kafitz et al., 1999; Kirn et al., 1994; Nottebohm, 1981; Thompson and Brenowitz, 2005; Vellema et al., 2014). Furthermore, many songbird species are sexually dimorphic in song behavior. In many species, such as the canary (*Serinus canaria*), only males sing. Female canaries rarely sing spontaneously, and never produce high quality songs under natural conditions (Pesch and Güttinger, 1985). However, these naturally non-singing birds can be persuaded to sing when treated with testosterone, as well as by a combination of both its active metabolites, DHT and estradiol (Devoogd and Nottebohm, 1981). Moreover, testosterone-induced changes in female canaries also include the masculinization of the song system (Nottebohm, 1980).

In a previous study we let female canaries develop their vocal motor skills by long-term implantation with testosterone, followed by implant removal to cease song production, and then by a second testosterone treatment to investigate the re-development of vocal motor skills (Vellema et al., 2019). We found that these birds showed a slow development of song features in the first treatment, with a faster re-acquisition during the second treatment. Furthermore, we provided evidence that the accelerated re-acquisition of vocal motor performance could be mediated by a lasting reorganization of the song motor circuit. Testosterone-induced singing led to substantial dendritic spine pruning in brain nucleus HVC of the song system, which was maintained after testosterone removal, whereas dendritic spine densities in another song premotor nuclei, nucleus RA, remained unaffected.

Here we demonstrate that unlike testosterone, DHT treatment of naive female canaries does not affect dendritic spine densities in nucleus HVC, and is insufficient to stimulate the development of song. However, once birds acquired singing skills through testosterone treatment, but stopped singing for some time, DHT alone was enough to stimulate the re-acquisition of previously acquired levels of song performance. These findings suggest that a significant shift in the hormone regulation of song production takes place during song learning, simplifying the mechanisms behind the re-acquisition of vocal motor performance. Such a simplified regulation of hormone-dependent vocal motor acquisition in experienced animals may play an important function in the ability to quickly recover previously acquired levels of vocal performance.

## Results

### Singing experience leads to a shift in hormonal modulation of song

We administered subcutaneous implants filled with the steroid hormone DHT (5α-dihydrotestosterone) to both naïve adult female canaries, who had never previously experienced neither hormonal treatment nor singing, and experienced females that had prior testosterone-induced singing practice, while recording their vocal output (figure 1A).

**Figure 1:**
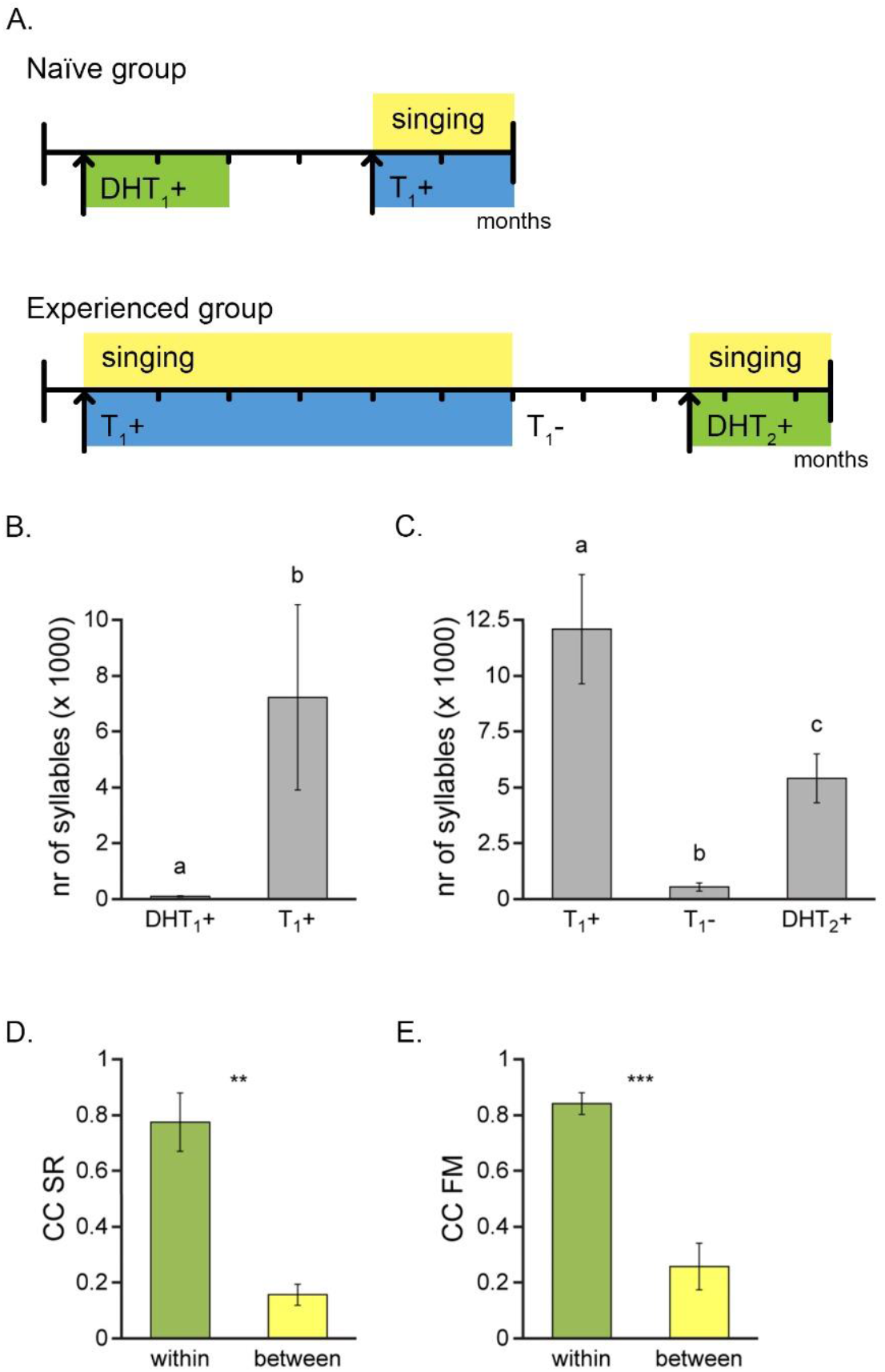
DHT triggers song development only after prior testosterone-induced singing experience. **(A)** Experimental setup. The naïve group was treated with DHT (DHT_1_+) for 2 months, which failed to trigger song production, and subsequently treated with testosterone (T_1_+) to confirm the singing ability of the individuals in this group. The experienced group was first treated with testosterone (T_1_+) and allowed to practice singing for 6 months, after which testosterone was removed and song production abolished for 2.5 months (T_1_-), followed by DHT implantation for 2 months (DHT_2_+), which eliciting singing. **(B)** DHT treatment of naïve female canaries (DHT_1_+) did not induce song production, while subsequent testosterone implantation (T_1_+) of the same individuals significantly increased the daily number of produced song syllables. **(C)** Another group of birds was stimulated to sing with testosterone (T_1_+), and ceased singing after testosterone withdrawal (T_1_-). Subsequent DHT treatment (DHT_2_+) of these individuals significantly increased the daily number of produced song syllables. **(D and E)** Similarity in song patterns after subsequent hormone treatments. Both the distribution of syllable rates (SR, D) and frequency modulations (FM, E) correlated strongly between T_1_+ and DHT_2_+ within individuals (green bars), and correlation coefficients (CC) were significantly higher than when cross-correlated between individuals (yellow bars). Columns represent the mean ± SEM (a, b, c: P < 0.05, Wilcoxon signed-rank test, n = 5 for B and n = 6 for C; ** P < 0.01, *** P ≤ 0.001, paired t-test; n = 6 for D and E).

In line with previous reports (Devoogd and Nottebohm, 1981), we show that treating naïve female canaries with DHT alone fails to stimulate singing (figure 1B). Subsequent testosterone treatment did, however, induce singing in the same individuals, confirming that the absence of song development during DHT treatment is not caused by an inability to sing, but is due to the ineffectiveness of DHT to stimulate song production in naïve animals. Surprisingly, DHT treatment of experienced female canaries that were previously allowed to undergo testosterone-induced song development did, on the other hand, succeed in stimulating singing (figure 1C).

Additionally, the observed song patterns that emerged during DHT treatment were very similar to the song patterns that developed during the preceding testosterone treatment (figure 1D and E). Both the distribution of syllable rates (SR) and frequency modulations (FM) correlated strongly between the first testosterone treatment and the subsequent DHT treatment within individuals (correlation coefficient, CC: 0.78 ± 0.10 for SR, and 0.84 ± 0.04 for FM), with significantly higher CC’s than when cross-correlated between individuals (P < 0.01, paired t-test). These findings indicate that DHT treatment can stimulate the development of song of similar quality as testosterone treatment, but only in birds that had previous testosterone-induced singing experience.

It is known that either testosterone, or a combination of both estradiol and DHT, is required to induce singing in naïve female canaries (Devoogd and Nottebohm, 1981; this publication), or in castrated male canaries (Sartor et al., 2005), suggesting that the activation of both AR and ER is needed to trigger song output. Our findings indicate that after an initial hormone-induced singing experience, the hormonal cascades modulating singing behavior are simplified, making AR receptor activation sufficient to trigger singing.

### DHT-independent synaptic pruning

In our previous study we found that testosterone-induced singing experience led to pronounced dendritic spine pruning in HVC, with dendritic spine densities remaining low as testosterone was removed and song production ceased (Vellema et al., 2019). Whether the reduction in spines is driven by the behavioral activation of the song motor circuit or whether spine pruning is directly induced by activation of the AR-pathway in HVC has remained elusive. By comparing dendritic spine densities in Golgi-stained HVC tissue of DHT-implanted naïve (DHT_1_+) and experienced female canaries (DHT_2_+) we found that, contrary to testosterone treatment, DHT treatment of naïve females did not lead to synaptic pruning in HVC (figure 2A). Both groups lacking singing experience (C and DHT_1_+) showed similarly high dendritic spine densities in HVC. On the other hand, singing DHT-treated experienced birds (DHT_2_+) had significantly lower HVC dendritic spine densities, similar to the other two treatment groups who experienced hormone-induced song development (T_1_+ and T_1_-; figure 2A).

**Figure 2.**
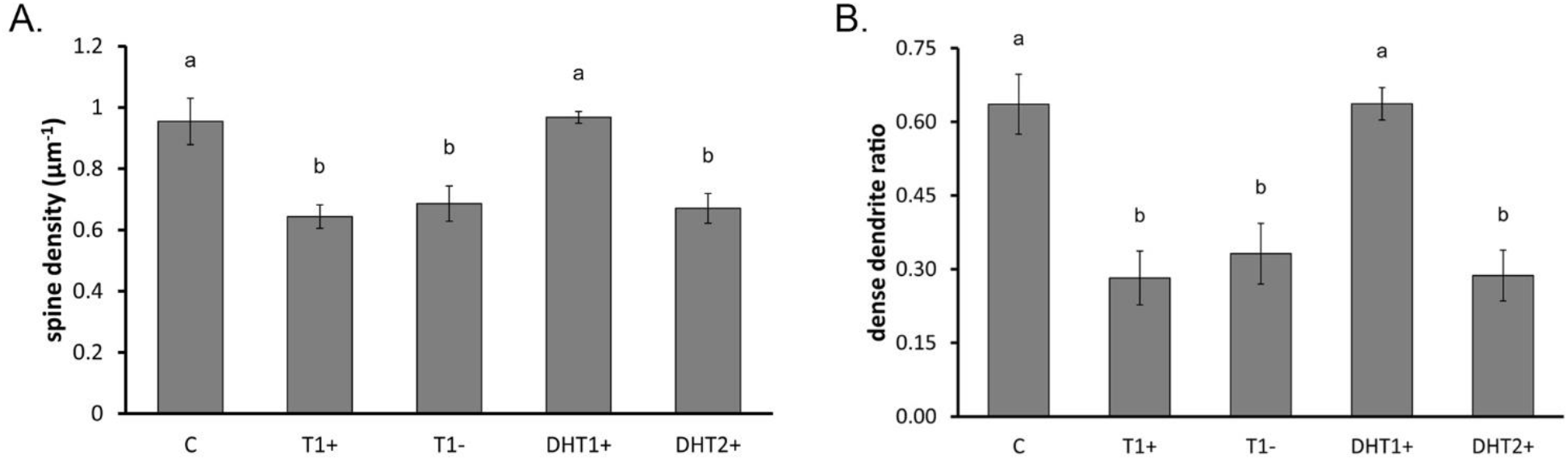
The singing-induced synaptic reorganization of HVC is independent of AR activation. (A) Compared to non-singing untreated control birds (C), spine densities were significantly reduced in testosterone-treated, singing birds (T_1_+), but not in non-singing DHT-treated naïve females (DHT_1_+). Spine densities remained significantly reduced up to 2.5 months after birds stopped singing by withdrawing testosterone (T_1_-), as well as after subsequent DHT-induced singing practice for 2 months (DHT_2_+). (B) Dense dendrite (> 0.8 spines per μm of dendrite) ratios further illustrate significant similarities between naïve groups (C and DHT_1_+) and birds who experienced singing practice (T_1_+, T_1_-, and DHT_2_+). [a, b: P < 0.01, ANOVA, n=6 animals. Data for groups C, T_1_+, and T_1_-taken from Vellema et al. (2019) with permission.]

After testosterone treatment, we previously found that the strongest reduction in spines was detected in the range of dendrites with the highest spine density (Vellema et al., 2019). Here we calculated the probability distribution of HVC dendrite types for DHT-treated females, and confirm that the changes in HVC’s synaptic circuit seems to mainly concern highly dense neurites. The ratio of dense dendrites in non-singing DHT-implanted naïve (DHT_1_+) females is similar to that of non-singing controls (C), whereas the ratio of dense dendrites is lower in all three groups that experienced testosterone-induced singing (T_1_+, T_1_-, and DHT_2_+; figure 2B). These densely spiny neurites predominantly belong to putative HVC_X_ neurons (Kornfeld et al., 2017), the permanent subpopulation of HVC projection neurons that are part of the pallial-basal ganglia-thalamic loop essential to vocal learning (Doupe et al., 2004).

Our results suggest that the observed changes in the HVC circuit are brought about by testosterone-induced activation of the vocal motor circuit, requiring more than AR activation alone. Furthermore, once consolidated, the HVC circuitry seems to be resistant to both testosterone removal and DHT treatment, as well as to singing practice offset or onset.

## Discussion

In this study we show that the reorganization of the song motor circuit evoked by singing practice is accompanied by a shift in the hormonal cascades needed to trigger singing. The testosterone-induced practice of vocal motor skills lead to both a simplification of the song connectome by the elimination of superfluous synapses, as well as to a simplification of the sex hormone-modulated pathways required to elicit song behavior.

Both ER activation by estradiol as well as AR activation, by either testosterone or its more potent metabolite DHT, are required to trigger song acquisition in female canaries (DeVoogd and Nottebohm, 1981). The necessity for ER activation is supported by studies showing that active brain aromatase, the enzyme responsible for the synthesis of estradiol from testosterone, is needed to induce singing in testosterone-treated naïve females (Brenowitz and Lent, 2002, 2001; Fusani et al., 2003). However, here we show that if birds learned to sing once before, but stopped singing for some time, ER activation is no longer required and DHT alone is sufficient to induce the re-acquisition of song.

The mechanisms and gene cascades involved in this complex interplay between hormonal and behavioral effects on brain circuits are not fully understood. Previous studies revealed that testosterone-induced gene expression in HVC is related to cellular differentiation, axon, dendrite and synapse organization (Dittrich et al., 2014; Frankl-Vilches et al., 2015). Furthermore, whereas AR have been found in many song system nuclei, including both HVC and RA, ER have only been found in HVC (Gahr et al., 1987). This makes HVC the only song nucleus expressing both AR and ER, and many active genes in HVC contain androgen and/or estrogen response elements (Frankl-Vilches et al., 2015).

AR and ER are expressed in distinct cell populations within HVC (Gahr, 1990a), and ER are mainly restricted to HVC_X_ projecting neurons (Gahr, 1990b). Interestingly, HVC_X_ neurons are also the neuron population that is most strongly affected by spine pruning upon testosterone-induced singing (figure 2B). Estrogens have been shown to affect dendritic spine density and induce the remodeling of synapse structure and function in mammals (Luine et al., 2018; Srivastava, 2012), as well as facilitate memory consolidation (Frick, 2015). Perhaps ER activation is responsible for promoting synaptic plasticity in HVC_X_ neurons in a first singing experience, leading to the optimization of the motor circuit for song production. Once circuit optimization is accomplished, motor memories are stored in the responsible synaptic circuits, and the ER-activated gene cascades that promote synaptic plasticity become unnecessary or even detrimental for the preservation of these optimized circuits. Thereafter, ER-activated gene cascades may thus permanently be switched off in order to preserve the stored motor memories. Future investigations should explore gene expression across different cell populations in HVC to determine the experience-dependent gene cascades involved in the hormonal regulation of song behavior.

Similar to what we observed in female canaries, there may also be a differential role for AR and ER activation during song development in naturally-raised male canaries. Although juvenile males express high levels of both receptors in HVC, AR-expression persists throughout the breeding season, while ER-expression is significantly down-regulated (Gahr and Garcia-Segura, 1996; Gahr and Metzdorf, 1997). Thus in both male and female canaries, AR and ER signaling pathways are both necessary during song ontogeny, but once vocal skills have been acquired, AR pathways appear to play the more dominant role in the maintenance and re-acquisition of song performance.

Extensive evidence exists for sex hormones’ influence on both brain plasticity, as well as language development, cognition, learning, and memory (Anthoni et al., 2012; Bailey et al., 2017; Frick et al., 2015; Friederici et al., 2008; Hahn et al., 2016; Hamson et al., 2016; Hawley et al., 2013; Leonard and Winsauer, 2011; Postma et al., 2000; Schaadt et al., 2015; Vahaba and Remage-Healey, 2018). Nonetheless, there are still gaps in our understanding of the complex interplay between sex hormones, brain, and behavior. Closing these knowledge gaps might help explain the contradictory results obtained in the use of sex hormones for the treatment of cognitive declines and psychiatric disorders (Beer et al., 2006; Cherrier et al., 2007; Elias and Kumar, 2007; Resnick et al., 2017; Wolf et al., 2000). Our findings provide a further piece of this incomplete puzzle. Understanding the hormone pathways and gene cascades involved in the formation, maintenance, and re-acquisition of vocal motor skills in our female canary model system has the potential to both further our knowledge on motor learning and motor skill savings, as well as to inspire better treatments for disorders affecting learning and memory.

## Materials and Methods

### Subjects

For this study, one-year old female domesticated canaries (*Serinus canaria*) were taken from the breeding colony of the Max Planck Institute for Ornithology in Seewiesen, Germany. Experimental procedures were conducted according to the guidelines of the Federation of European Animal Science Associations (FELASA) and approved by the council for animal experimentation of the Danish ministry of environment and food.

### Hormone treatment

In the DHT control group (DHT_1_+), birds were subcutaneously implanted with 8 mm silastic tubes (Dow Corning, Midland, MI; ID: 1.47 mm) filled with 5α-dihydrotestosterone (DHT). After 2 months of DHT implantation, birds were either sacrificed in order to collect brains for Golgi-Cox processing (n=5), or DHT implants were removed for subsequent singing ability confirmation, by subcutaneous implantation with 8 mm silastic tubes filled with testosterone (n=5). In the DHT re-treatment group (DHT_2_+, n=6), birds were subcutaneously implanted with 8 mm silastic tubes filled with testosterone, which were removed after consolidation of song, approximately 6 months later. After 2.5 months in which birds did not produce song, a second 8 mm silastic tube containing DHT was implanted, and birds were sacrificed 2 months later.

### Hormone analysis

Throughout the experiments, blood samples were collected from the birds’ right wing veins using heparinized hematocrit capillaries (Brand, Wertheim, Germany). Directly after blood collection, samples were centrifuged at 3000 RPM for 10 min to separate cells from plasma, and stored at −80 ºC, until analysis. Testosterone and DHT concentrations in the blood plasma were determined by radioimmunoassay (RIA) after extraction and partial purification on diatomaceous earth (glycol) columns, following the procedures described in (Goymann et al., 2001; Goymann et al., 2006). Plasma levels of both testosterone and DHT during the different hormone treatments are shown in Table 1.

**Table 1:**
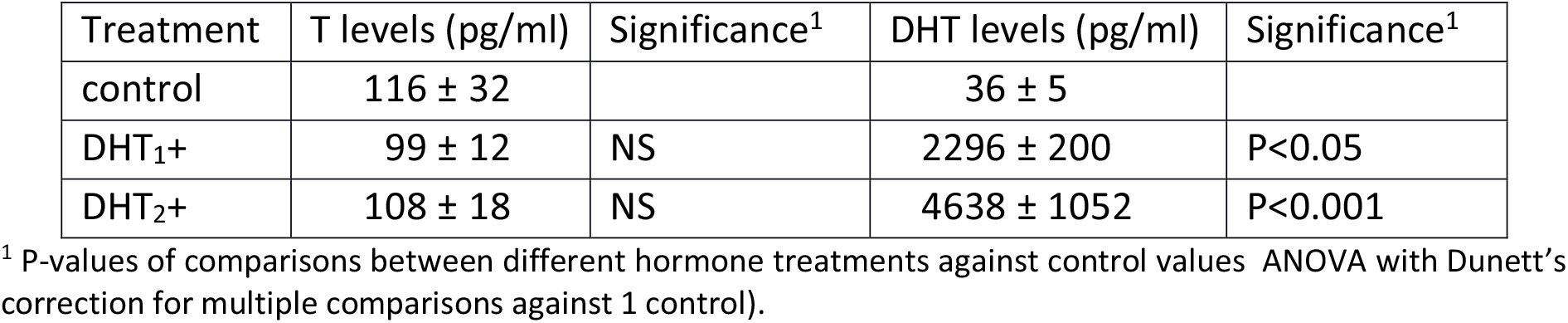
Plasma levels of testosterone (T) and 5α-dihydrotestosterone (DHT) during different hormone treatments.

### Song recordings and analysis

Birds were kept in sound attenuating chambers and the vocal activity of each bird was recorded using Sound Analysis Pro 2.0 (SAP, Tchernichovski et al., 2004). Vocalizations were segmented into individual syllables with the fully automated Feature Batch module in SAP by applying an amplitude threshold to the sound wave. To filter out non-song vocalizations as much as possible, syllables were only included when produced within a song bout of at least 750 ms. A song bout was defined as a sequence of sounds traversing the amplitude threshold with an interval of no more than 100 ms. For each syllable, frequency modulation (FM), duration (d), inter-syllable interval (i), and syllable rate (SR) were calculated and stored in MySQL 5.1 tables (Oracle, Redwood Shores, CA; http://www.mysql.com). The syllable rate was defined for each syllable as: SR_a_ = (d_a_+i_a_)^-1^, where d_a_ is the duration of syllable a, and i_a_ is the interval between the end of syllable a and the beginning of the consecutive syllable within the same song bout, irrespective of syllable type.

To investigate the dynamic change of song features over the course of the experiment, daily histograms were obtained by rounding individual SR or FM values to the nearest integer, and plotting the number of times those integers occurred each day. Developmental correlation plots for both SR and FM were produced by calculating the Pearson product-moment correlation coefficient (CC) between the average daily histogram of 1 week at the time of song stabilization and all other recorded days. A non-linear exponential curve (1) was fitted through the CC values and song parameters were considered stable as soon as the daily increase in similarity dropped below 0.001 CC/day.

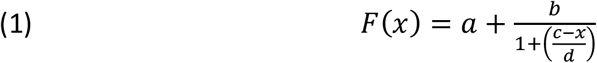

### Spine and dendrite quantification

Brains were collected immediately after birds were sacrificed and processed for Golgi-Cox staining using the FD Rapid Golgi Stain kit (FD NeuroTechnologies, Columbia, MD) as previously described in Vellema et al. (2019), in order to produce Golgi-stained brain sections for spine quantifications. For each bird, z-stacks of nucleus HVC were obtained from three brain sections with a Nikon Eclipse Ti microscope (Nikon, Tokyo, Japan), equipped with a 60x oil immersion lens (CFI Plan Apo VC 60x Oil). The outline of each HVC section was delineated in ImageJ (http://rsb.info.nih.gov/ij/), and 100 μm^2^ non-overlapping regions of interests (ROIs) were randomly placed within the boundaries of HVC prior to the quantifications. Quantifications were carried out as previously described in Vellema et al. (2019). All quantifications were conducted by an experimenter that was blind to the experimental condition of the animals.

### Statistical analysis

To determine differences in hormone levels, data from all experiments were pooled and plasma values during the different hormone treatments were compared using a one-way ANOVA with Dunett’s correction for multiple comparisons against one control value (untreated birds). The average daily syllable production was compared using a Friedman test, followed by a Wilcoxon signed-rank test to compare time points.

Song pattern similarities during the 1^st^ and 2^nd^ hormone treatments were established by calculating Pearson’s CC between the 7-day average of the SR or FM patterns from stabilized song during the 1^st^ treatment with the 7-day pattern average from stabilized song during the 2^nd^ treatment, constituting the within-individual CC. Between-individual CCs were determined by cross-correlating the 7-day pattern average during the 1^st^ hormone treatment of each bird with the 7-day pattern average during the 2^nd^ hormone treatment of each of the other birds. Differences between within-individual CCs and between-individual CCs were compared using paired-samples t-tests.

All statistical tests were two-tailed, and considered significant if the coincidence interval of 95% was exceeded. Measurement values in the text are given as means ± SEM, unless stated otherwise.

## Acknowledgements

The authors would like to thank Ingrid Schwabl, Monika Trappschuh and Wolfgang Goymann for performing and analyzing the hormone radioimmunoassays. This work was supported by the European Union’s Horizon 2020 (H2020) Marie Skłodowska-Curie grant nr. 701660 to MV, and the Max-Planck-Gesellschaft (MPG) to MR and MG.

